# The Miniature-chemostat Aging Device: A new experimental platform facilitates assessment of the transcriptional and chromatin landscapes of replicatively aging yeast

**DOI:** 10.1101/363523

**Authors:** David G. Hendrickson, Ilya Soifer, Bernd J. Wranik, Griffin Kim, Michael Robles, Patrick A. Gibney, R. Scott McIsaac

## Abstract

Replicative aging of *Saccharomyces cerevisiae* is an established model system for eukaryotic cellular aging. A major limitation in yeast lifespan studies has been the difficulty of separating old cells from young cells in large quantities for in-depth comparative analyses. We engineered a new platform, the Miniature-chemostat Aging Device (MAD), that enables purification of aged cells at sufficient quantities to enable genomic and biochemical characterization of aging yeast populations. Using the MAD platform, we measured DNA accessibility (ATAC-Seq) and gene expression (RNA-Seq) changes in aging cells. Our data highlight an intimate connection between aging, growth rate, and stress, as many (but not all) genes that change with age have altered expression in cells that are subjected to stress. Stress-independent genes that change with age are highly enriched for targets of the signal recognition particle (SRP). By obtaining pure populations of old cells, we find that nucleosome occupancy does not change significantly with age; however, significant age-dependent changes in accessibility at ~12% of genomic loci reflect decreased replication and changing activities of cell cycle and metabolic regulators. Finally, ATAC-seq revealed that upregulating the proteasome by deleting *UBR2* reduces rDNA instability usually observed in aging cells, demonstrating a connection between proteasome activity and genomic stability.

## Introduction

Aging is a multifactorial process characterized by loss of homeostasis, reduced fitness, and increased risk of death. In people, aging is the primary risk factor for a myriad of ailments including dementia, cancer, and heart disease [1]. Across different organisms, aging is influenced by both environmental [2–4] and genetic factors [5]; defining a quantitative systems-level understanding of the molecular changes that cause aging remains an open problem.

The unicellular eukaryote *Saccharomyces cerevisiae* (budding yeast), which typically produces ~25 daughter cells before dying [6], has enabled several insights into the cellular aging process via a combination of cell biological, genetic, and genomic approaches [7–11]. For example, instability at the ribosomal DNA (rDNA) locus, which contains ~100-200 tandemly repeated copies in the genome, can result in the formation of rDNA extra-chromosomal circles (ERCs), and is is a major determinant of yeast replicative lifespan (RLS) [11–15]. A single nucleotide polymorphism that reduces the rate of origin firing at the rDNA locus increases rDNA stability, and thereby increases RLS [16,17]. In a seemingly unrelated pathway, a decrease in vacuolar acidity early in a mother cell’s life promotes mitochondrial dysfunction and limits her lifespan [7]. Likewise, caloric restriction-mediated life extension, acting through the homeostatic regulator *GCN4*, is a well-studied yeast RLS intervention that is broadly phenocopied by perturbations that inhibit translation [18,19]. In a step towards a systems-level understanding the genetic determinants of cellular aging, RLSs of >4,500 gene deletions were recently measured using microdissection to physically separate daughter cells from mother cells and manually counting the number of divisions before mother cell death [8]. Even still, no unifying model has coalesced from these disparate observations. The rarity of old mother cells in mitotically growing populations and the large number of cells required for genomic and biochemical assays has hampered our ability to dynamically monitor genomic changes in aging cells that would aid in building and testing explanatory models of aging.

With currently available technology, obtaining large numbers of old mother cells requires physical separation by magnetic cell sorting, which can be combined with daughter cell-killing genetic programs to increase the average age and purity of purified populations [20,21]. Profiling aging population*s densely* across multiple ages or broadly across genotype and environment can be improved with fluidic approaches [9]. With MAD, the fluidic setup is straightforward, no genetic engineering of strains is required and the cells are aged in a near-constant environment with fresh media continuously provided. To our knowledge, no genomic assay (RNA-seq, ATAC-seq, etc.) has been published on strains harboring longevity-associated mutations as they age. Identifying molecular changes, and perhaps ultimately distinguishing different ‘trajectories’ cells take as they age, would be improved with new methods that enable rapid purification of large quantities of replicatively aged cell populations.

Here we introduce a robust methodology, utilizing miniature chemostats [22], for purifying populations of aged yeast cells that is both easily scalable and highly flexible with respect to environmental condition and strain background. Applying next generation sequencing-based approaches, we profile chromatin and transcriptome changes (using ATAC-Seq and RNA-Seq) in wild type strains and strains harboring longevity-associated mutations at different replicative ages. Comparing our data to the literature we were able to verify some observations of aging cells, falsify a prominent dogma, and identify several features specific to aging cells. Consistent with previous findings, we observe a clear connection between aging, slow growth, and stress. Contrary to previous findings, we show that a global increase in DNA accessibility is not a general feature of aged cells. We explore the relationship between accessibility and instability at 2 the rDNA locus and find a novel connection between proteasome activity and regulation of rDNA and nucleolar morphology with age. Collectively, despite differences at the rDNA locus, age-related changes in gene expression and chromatin are strikingly similar across longevity mutants.

## Results

### Ministats: A method for the enrichment of large numbers of replicatively aged yeast cells

To collect pure populations of aged mother cells, we first labeled the cell walls of exponentially growing cells with biotin and grew them ~12 hours overnight in liquid cultures before capturing biotinylated cells with magnetic streptavidin beads, as previously described [23] (Fig. 1A). Pre-growth in liquid culture after the biotin labeling improved the uniformity of the number of beads per cell, as compared to cells that were conjugated to beads immediately following biotin labeling. Following bead binding, the labeled mothers were loaded into our ministat culture vessel (~30 mL volume) surrounded by custom-sized neodymium ring magnets (Fig. 1A). Peristaltic pumps were employed to provide aging cells with fresh media flow and to wash away progeny. The optical density (OD _600_) of the culture media within the ministats was kept at less than 0.15 to ensure that pH and glucose remained constant in the vessels (data not shown). Mother cells were harvested by first removing the ministat culture vessels from the magnets and then washing cells in growth media to remove contaminating daughter cells. We verified that purified mothers had aged at the expected ~2 hour division rate in YNB glucose minimal media by counting the number of replication bud scars for both haploid and diploid strains (38 hours: haploid=15.5, diploid= 17.6; expected ~ 19) (Fig. 1B & C).

**Figure 1:**
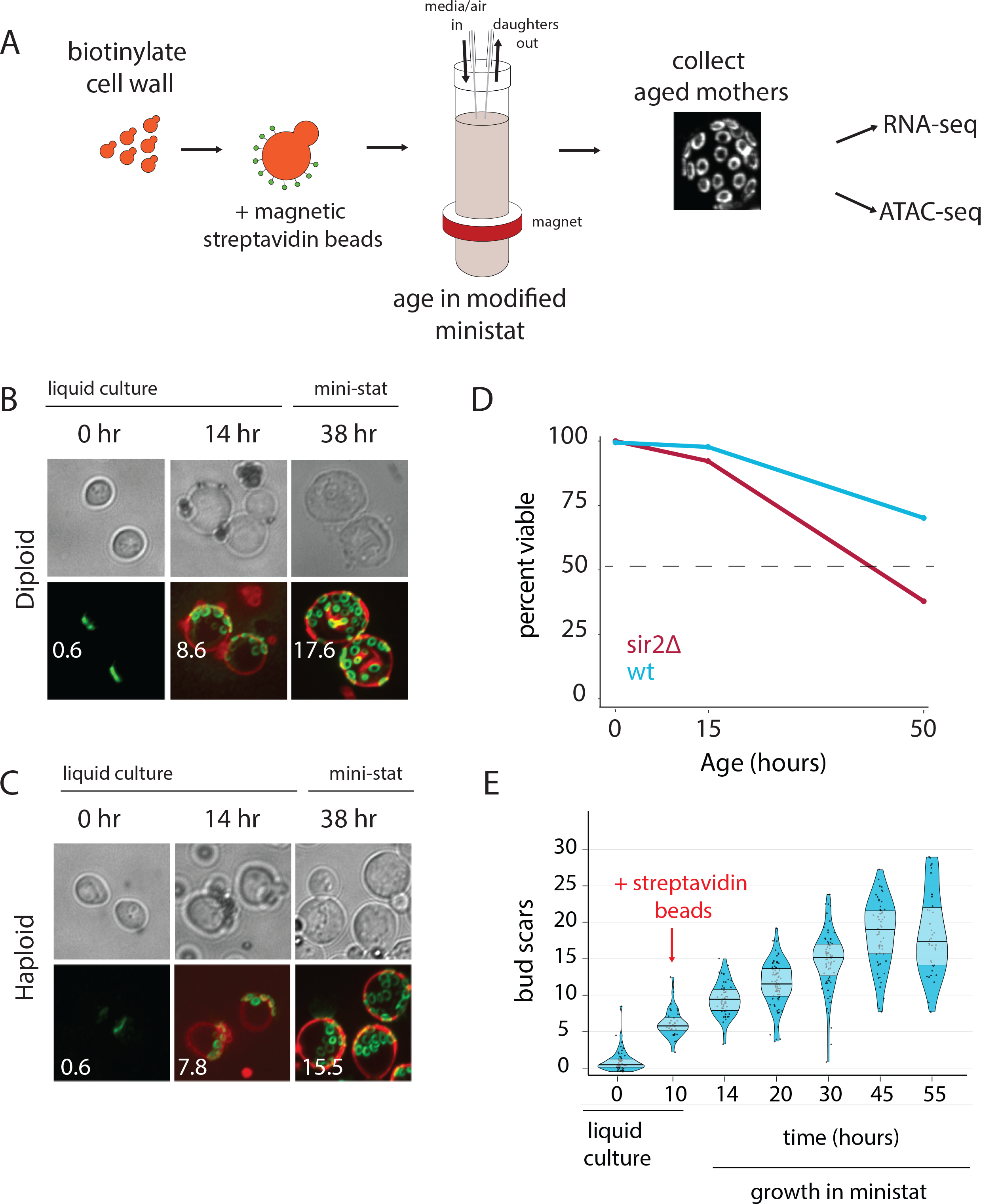
Overview of the MAD platform. (A) Yeast cells are biotylinated, grown in flask culture, labeled with streptavidin beads, and loaded into a modified miniature chemostats surrounded by neodymium magnets. Daughters don’t inherit the magnetic beads and are perfused from the culture, leaving behind a pure replicatively aged population of cells. (B,C). Haploid (DBY12000) and diploid (DBY12007) cells purified using MAD. (D) Viability of WT and *sir2∆* cells over a 50-hour aging time course. (E) Mother cell ages (bud scars) during 55-hour MAD time course.

Previous studies have used mother cell viability over time as a method for approximating differences in strain replicative lifespan (RLS) [20]. To test if we could observe differential aging of strains, we loaded separate ministats with wild type (WT) cells or short-lived *sir2*∆ cells, and measured membrane integrity as a proxy for viability with propidium iodide (PI) staining at three time points (Fig. 1D) [11]. The deletion of the *SIR2* NAD-dependent histone deacetylase shortens lifespan by promoting hyper-recombination at the rDNA locus [24,25]. As expected, viability for aged *sir2∆* mothers is ~2-fold lower than that of aged WT mothers (Figure 1D).

### Mother cells aged in the ministats exhibit hallmarks of the aging yeast transcriptome

Substantial previous work has established that aging yeast cells share some common features across different methods and strains [26]. As a starting point, we sought to validate the MAD method by testing if we could recapitulate the salient features of the aging yeast transcriptome while simultaneously uncovering novel age-dependent features by leveraging the ministat scalability to collect purified mothers over a dense aging time course. Thus, we collected aged cells from five separate ministats loaded from the same pre-culture (Fig. 1E) for profiling the aging transcriptome (RNA-seq) at multiple time points during yeast replicative aging. Age, viability, and purity (% of recovered cells that are mothers) were measured with microscopy at 3 each time point and were consistent with expectation from orthogonal methods (Fig. 1E, Table S1) [9,20].

We then used Sleuth (an RNA-Seq analysis software suite) to identify transcripts that change significantly with replicative age [27]. The significances (q-values) of these were then plotted against the magnitudes of age-dependent expression changes (Fig. S1A,B, Data S1) [27]. It is worth noting that we used an extensive catalog of *full-length* yeast transcripts (ORFs and non-coding RNAs, Table S2, [28–36]), including 5’ and 3’ UTR regions, for mapping sequencing reads. Strikingly, some of the most up-regulated transcripts were those arising from the rDNA locus, specifically from the Sir2p-silenced non-transcribed regions (NTS1 and NTS2). Transcription at the Non-Transcribed Sequence (NTS) locus is an aberrant Pol II event (almost undetectable in log phase culture) that has been previously shown to increase with age and ERC accumulation [37–39].

Next, we analyzed gene sets that are differentially expressed as a function of increasing age for enrichment of a variety of functional annotations using the Database for Annotation, Visualization and Integrated Discovery (DAVID; [40]). We found numerous significantly enriched terms related to metabolism, oxidoreductase activity, stress, and mitochondrial proteins amongst the genes whose expression increases with age (Data S1). Genes downregulated with age were enriched for multiple terms, including ribosome biogenesis, rRNA processing and translation as well as a host of other terms related to general cellular anabolic function (tRNA synthesis, amino acid biosynthesis, mRNA transport, exosome activity) (Data S1). In general, these results are consistent with previous reports that the aging transcriptome shares a high degree of similarity to that of induction of the environmental stress response (ESR) [9,10,41–43]. The ESR is the simultaneous activation of a set of stress-responsive pathways coincident with the down-regulation of ribosomal biogenesis and it is inextricably tied to growth rate (i.e. if a yeast culture is growing slowly, the ESR is up-regulated) [44]. Likewise, we also observed a shift away from glycolysis towards gluconeogenesis, similar to previous reports [45,46]. Thus, MAD robustly captures the transcriptomic hallmarks of yeast aging that have been reported across a diversity of strains and methodologies [26].

### The ESR is a robust feature of the aging transcriptome

We attempted to resolve the earliest age-dependent transcript changes using hierarchical clustering of genes with significant age-dependent expression (q < 0.05) (Fig. 2). We found that age-dependent transcripts display a clear temporal progression as we observed clusters composed of early-, middle-, and late-age changes. To understand what processes and events these clusters might be representative of, we included transcript annotation categories alongside the heatmap for multiple classes of non-coding transcripts, sub-telomeric genes, TY retrotransposon genes, and ESR-induced and ESR-repressed genes (Fig. 2, Table S2).

**Figure 2:**
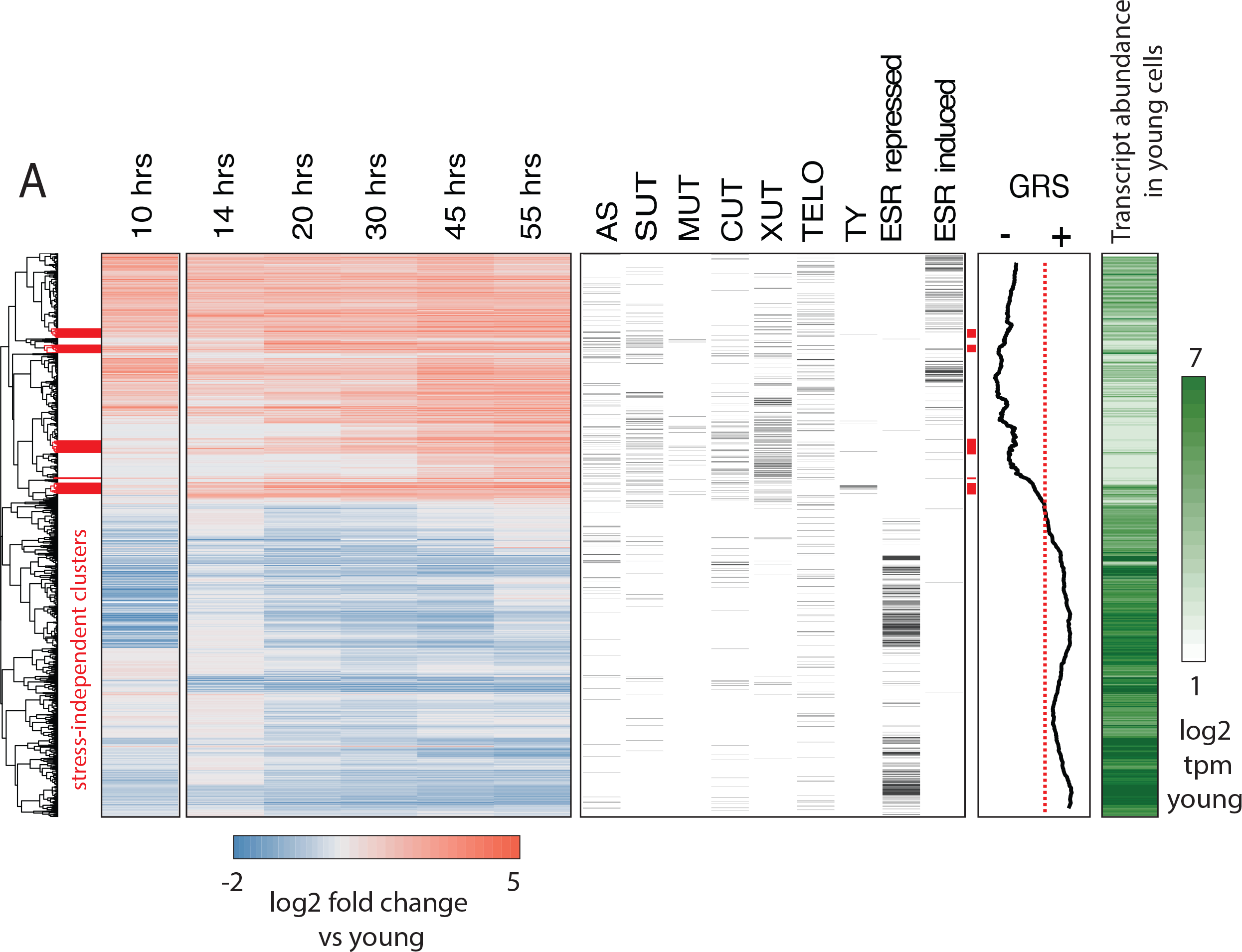
Age-dependent gene expression dynamics. (left) Hierarchically-clustered heatmap of transcriptome at increasing replicative ages ~2900 genes (the transcriptome at 10 hours is conflated with cell beading). Different transcript types are labeled in black directly to the right of the transcriptional data. (AS = antisense, SUT = stable unannotated transcript, MUT = meiotic unannotated transcript, CUT = cryptic unstable transcript, XUT = Xrn1-sensitive unstable transcript, TELO = subtelomeric, TY = TY repeat elements, ESR = Environmental stress response). (right) Moving window average of (100 gene bins) of Growth Rate Slopes from *Brauer et al*. and log2 transcript abundances in transcripts per million during initial log-phase growth (green). Red bars indicate clusters displaying early-age induction independent of the ESR.

We found that an unmistakable feature of the aging transcriptome is the activation of the ESR (i.e., almost all ESR-induced genes were upregulated with age and almost all ESR-repressed 4 genes were down-regulated [Fig. 2]). We also observed an age-independent initial rise and decline prior to the age-dependent increase in ESR induction in response to the beading process at the 10 hour time point. Subsequent iterations of the protocol were modified to reduce beading stress (Supplementary Note); importantly, age-dependent changes and ESR induction from both protocols were highly correlated confirming that our observations are reflective of yeast aging in general.

In order to extend our comparison between the ESR and the aging transcriptome, we used Growth Rate Slopes (GRS) from Brauer *et. al*. [44]. By modulating the growth rate using nutrient limitation and measuring global patterns of gene expression across a range of growth rate, Brauer *et. al*. characterized the magnitude/direction of the effect of growth rate on gene expression as the *slope* of the regression of gene expression on growth rate [44]. Gene-specific GRSs have a straightforward biological interpretation. Genes with negative GRSs increase expression as growth rate slows, while genes with positive GRSs decrease expression as growth slows. We plotted the moving average for 100 gene bins across the heatmap in Figure 2 and found that the remarkable relationship between age-related change and stress-related change extends to almost all genes for which a GRS was measured.

### Dense profiling of aging yeast cells reveals early transcriptomic events independent of the ESR

Our data and the literature agree that the aging yeast cell experiences stress/slowed growth [47]; however the root cause[s] remain elusive. We reasoned that potentially causative events would occur both before and independently from the stress signature. Thus, we highlighted ESR-independent transcripts and processes that happen earliest in our time course as prospective initiating agents of yeast replicative aging. As such, we isolated and analyzed age-induced gene clusters that were relatively depleted of ESR induced genes (Fig. 2 red bars, Data S1). Strikingly, this group of genes was highly enriched for TY transposon genes (Fig. 2, Data S1). Although age-dependent increases in transposon expression have been previously reported [21], here we show that it is one of the earliest and strongest age-related changes (Fig. 2, Data S1). We also found multiple transcripts arising from the non-transcribed spacer region (NTS) of the rDNA repeat enriched within this gene set as well as genes known to be involved with response to replicative stress and DNA damage (*RNR3*, *IRC4*) raising the possibility that rDNA instability and Extrachromosomal rDNA circle (ERC) formation are very early events in the yeast aging process.

The carbon source-responsive zinc-finger transcription factor, *ADR1*, was among these genes and was previously reported to be an early-age transcriptional responder [43]. *ADR1* has also previously found to be involved in regulating gluconeogenesis during the diauxic shift, raising the possibility that *ADR1* has a role in regulating the age induced transcriptional shift towards gluconeogenesis [48]. In addition, these clusters contained regulators of oxidative stress that are induced with age, including *SRX1*, *SOD2*, *TSA2*, *GPX2* and *XBP1*. Together, these data revealed several compelling leads that are independent of the ESR for further study into the originating deleterious events during replicative aging in yeast.

### Low abundance ORFs (at young ages) and non-coding RNAs tend to increase with age

In general, we observed a negative correlation between transcript abundance during early age and age-related induction (i.e. low abundance transcripts tend to undergo the largest fold increases; Fig. S2). Interestingly, we also found that non-coding transcripts (Fig. 2, right panel) typically fell into the age-induced clusters similar to previous results [49]. A recent study from Hu *et al*. suggested that all transcripts increase with age as a result of pervasive global transcription stemming from nucleosome loss [10]. Consistent with this hypothesis we observed that many induced genes are either expressed at low or undetectable levels during exponential growth (Fig. 2, right bar, white to green). However, a universal increase in transcription with age *cannot* explain numerous induced genes that are highly expressed and functionally related. The tendency of low abundance and silenced transcripts to increase with age might be explained by the decrease in exosome components we observed, as well as a decline in mRNA degradation pathways (Fig. S1B) and decreasing transcriptional fidelity [49]. This is consistent with the observation that the exosome component XRN1-sensitive unstable transcripts (XUTs) and cryptic unstable transcripts (CUTs) tend to be age-induced (Fig. 2, Data S1). Lastly, we re-analyzed the spike-in normalized transcriptomic data from Hu *et al*., and found that the observed universal induction of gene expression is not consistent between replicates (Fig. S3).

### The ESR is highly correlated with the aging transcriptome across mutant panel

Although multiple wild type strains have been transcriptionally profiled during the aging process, to our knowledge, the aging transcriptome of mutants has not been studied. It is unknown if long- and short- lived strains experience the same process albeit on a different timescale or if there are clear differences in what stresses RLS mutants undergo as a function of age and how their response might differ as a function of genotype. In the event of the latter scenario, perhaps different long-lived strains share aspects unique from that of wild type such that a “protective” transcriptional response or state might be identified. To explore this idea further, we chose to profile two classically long-lived strains (*ubr2∆* and *fob1∆*) whose RLS phenotypes are reported to be independent. Deletion of *FOB1* extends yeast RLS through modulation of rDNA stability and ERC formation [11,50] whereas deleting *UBR2* could confer longevity through constitutive up-regulation of proteasome activity [51]. In addition, we selected the short-lived *sir2∆* strain to profile with age as well. As mentioned above, *SIR2* silences transcription of the NTS rDNA region and is a major regulator of rDNA stability.

We first determined that none of our mutant panel had significant growth rate defects (Fig. S4A) and could be grown in the MAD for equivalent amounts of time. We then grew and labeled mother cells as described above for growth in ministats and sampled cells at three timepoints: young (exponential growth phase), middle-aged cells (mothers with 20 hours total growth), and old cells (mothers 40 hours of total growth). We ran three separate aging campaigns separated by one week as independent biological replicates and prepared samples for RNA-seq and ATAC-seq as mentioned above. We assessed purity and viability of our samples using flow cytometry as previously reported and median age of mother populations via microscopy ([9], Fig. S5, Table S1).

We then checked that we could observe reported differences between strains (e.g., up-regulation of proteasome components in the *ubr2*∆ strain) (Figs. S4B, S4C). Importantly, we also observed a significantly lower median bud scar age at both aged time points for the *sir2*∆ strain compared to wild type and slightly higher ages for *ubr2∆* and *fob1∆* compared to wild type in both the total population of collected cells and in the viable fraction (Fig. S6A,B). That cell division slows in the approach to replicative senescence is a well-documented observation [52]. Therefore, we reasoned that the older bud scar age in *fob1∆* and *ubr2∆* is not a function of a faster growth rate per se, but rather a result of there being a higher percentage of actively dividing cells in *fob1*∆ and *ubr2*∆ backgrounds.

After quantifying transcript levels, we used Sleuth to call significant gene expression changes as a function of age. Notably, since we calculate the magnitude of change as an exponential fit of expression change to numbers of bud scars, our “aging slope” can be interpreted as rate of change per cell division. We then used the aging slope to determine both the strain-dependent and strain-independent changes dependent on increasing age (Fig. 3A, Fig. S7). Although the aging transcriptomes were similar, it was the striking paucity of significant differences that emphasizes how analogous the transcriptional aging profile appears across strains (Fig. S7). Even for *sir2*∆, the strain that exhibited the strongest strain-specific response to aging, the significant differences were in most cases a matter of magnitude per replication event (bud scar); the same genes changed, only more so, reflective either of a higher aging rate or a larger percentage of cells close to replicative senescence (Fig. S7).

**Figure 3:**
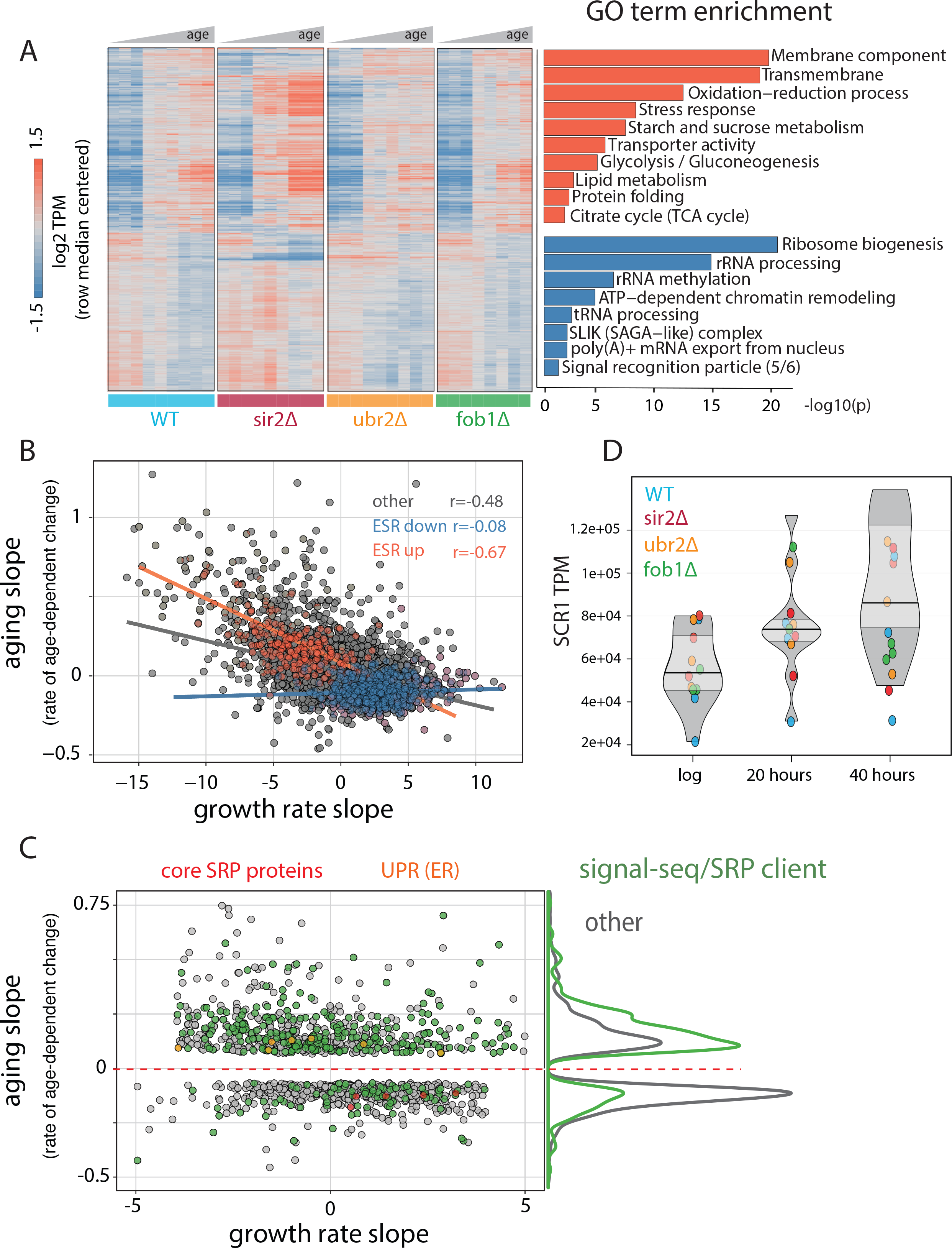
Common features of the aging transcriptome across multiple genotypes. (A) (left) Hierarchically-clustered heatmap of ~1500 genes with significant age-dependent transcriptional responses across multiple genotypes. (right) Significance of GO terms in up- and down- regulated genes, respectively (-log10(p)). (B) Scatterplot of “aging slopes” from (A) versus “growth rate slopes from Brauer *et al*. Canonical ESR-induced transcripts are labeled red and canonical ESR-repressed transcripts are labeled blue. (C) Subset of (B), filtered for genes with aging slopes q < 0.05 are and growth rate slopes > 4 or < −4 removed. SRP-encoding genes are labeled red, UPR genes are labeled orange, and SRP clients are labeled green. Right panel: density of SRP clients (green) and other genes (grey) across significant aging slopes. (D) Gene expression levels of *SCR1* at increasing ages.

To identify functionally coherent patterns across the significant changes common to all genotypes, we used GO term analysis to find enriched functional annotations for the set of genes that increase with age (Fig. 3A, red) and the set of genes that decrease with age (Fig. 3A, blue) across the tested genotypes. The enriched annotation sets for genes that increase with age across genotype mirror the terms we found for the aging wild type strain in our previous dense time course (e.g., stress, membrane component, oxidation-reduction processes) going up as a function of age and ribosome biogenesis and rRNA processing going down with age. Notably, we observed an age-dependent decrease in five out of six components components of the signal recognition particle (SRP).

We compared the rate of change with age (Fig. 3B, y-axis) to the rate of change with growth rate (Fig. 3B, x-axis) and found a strong correlation for genes that increase with age and canonical ESR induced genes (r = −0.67). In general, the correlation for all genes (r=-0.48) was strong and underscores the close relationship between RLS, stress, and growth rate.

### Age-dependent changes in gene expression that are independent of the ESR are highly enriched for SRP targets

As with the first WT time course, we sought to extricate age-dependent changes in gene expression that are not dependent on growth rate and the ESR. Accordingly, we focused on the set of significant age-related changes bounded by a GRS between −5 and 5, a regime wherein the age-dependent changes in gene expression that we observe are not clearly dominated by the apparent relationship between age-related change and GRS/ESR (Fig. 3C). We next asked whether this class of genes was a representative subset of all age-dependent changes genes or if it was enriched for specific functional annotation sets using GO term analysis (Data S2). We found that genes with an age-related increase in expression that are not strongly dependent on the GRS are greatly enriched for genes coding for transmembrane helices and more specifically for those that are classified as secreted and/or of containing sequence for a signal recognition peptide (SRP binding site) (Data S2). Next, we compiled a more exhaustive curated list of transcripts predicted or known to associate with the SRP (signal sequence containing and SRP-binding; [53], Data S2, Fig. 3C green dots). We found that these SRP clients are, as a class of genes, significantly upregulated with age in a GRS-independent manner (p < 10^−51^, hypergeometric, Fig. 3C). As SRP clients tended to increase in expression with age in parallel with age-dependent downregulation of SRP subunits (Fig. 3C, red dots), we asked if these two phenomena might be linked. To this end we plotted the expression of the SRP subunits versus the expression of SRP clients across yeast deleteome dataset, in which global gene expression was profiled across ~1500 deletion mutants ([54], Fig. S8). Expression of the SRP subunits and SRP clients were strongly negatively correlated (r ~ −0.5), suggesting the existence of a negative feedback loop between SRP and its clients.

Beyond its constituent proteins, the SRP complex also contains a non-coding RNA subunit, *SCR1*. Given that our RNA-seq libraries were depleted for rRNA rather than polyA-selected (*SCR1* is not polyadenylated) we were able to check the effect of age on *SCR1* levels. *SCR1* is a highly abundant transcript (5-10% of the non-ribosomal transcriptome) and highly structured, both features that increase noise in RNA-seq experiments. Even so, the aggregate trend across all of the genotypes we assayed indicates that *SCR1* is increasing roughly two-fold from ~5% to ~10% of the transcriptome. We also checked the abundance changes for *SCR1* in our dense time course and found the same upward (with age) trend (Fig. S9). As the central component/scaffold of the SRP, *SCR1* directly binds each protein subunit. Thus, we hypothesized that an increase in *SCR1* abundance coupled with a decrease in SRP protein expression might disrupt SRP complex stoichiometry and SRP-mediated translocation and translation efficiency. Consistently, genes involved in ER-associated protein degradation (ERAD), specifically the members of the Cdc48p-Npl4p-Ufd1p segregase complex, were upregulated with age (Fig. 3C, orange dots) suggesting that aging cells experience ER stress. Importantly, the SRP components, the SRP clients, and the ER unfolded protein response are not canonical ESR-related genes or genes with a strong GRS. Thus, the putative age induced malfunction in proper ER function cannot simply be explained as part of the normal yeast response to stress and/or slow growth and is detected in both the assayed short and long-lived strains. These results are consistent with reported role for ER stress in regulating yeast RLS [55,56].

### Proximity to telomeres correlates with age-dependent increases in gene expression

Clustering analysis revealed that genes positioned in subtelomeric regions are more likely to increase with age than to decrease (Fig. 2). In addition, previous work has suggested a loss of silencing at subtelomeric regions, potentially as a result of redistribution of *SIR* proteins from telomeres to the nucleolus, follows from an increasing rDNA copy number and age-dependent nucleolar dysfunction that is not dependent on telomere length [57–59]. We thus sought to determine to what extent telomere proximity affects the expression of a gene as a function of replicative age. To this end, we looked at rate of change with age for every gene versus the distance in base pairs from the telomeres for each genotype (Fig. S10A). We found that genes located in subtelomeric regions are more likely to increase in expression with age (Fig. S10A). Given the link we observed between increases in gene expression with age and increased expression as a function of slow growth and stress discussed above, we next asked if the observed age-dependent upregulation of sub-telomeric gene expression was also similarly connected. We plotted the GRS versus distance from the telomeres and found that indeed, subtelomeric genes as a group have a negative GRS and thus increase in expression as a function of slower growth rate (Fig. S10B).

### Global nucleosomal occupancy does not decrease with age

Having characterized the aging transcriptome across a panel of strains, we sought to thoroughly characterize the aging chromatin structure. ATAC-seq is an assay for measuring chromatin accessibility and nucleosomal occupancy and positioning by counting insertions of Tn5 transposase in the genome [60]. We used ATAC-seq to examine changes in the chromatin structure during yeast aging. We extracted transposase insertions from end points of ATAC-seq fragments and counted the number of insertions at each genomic location as a proxy of chromatin accessibility. To correct for variable sequencing depth and different amounts of rDNA between samples, insertion counts were normalized so that the total number of non-ribosomal, non-mitochondrial, uniquely mapped insertions was the same between samples. Precision of high-resolution analyses of chromatin structure are sensitive to read depth [61], and therefore we sequenced each sample deeply, requiring at least 30 million uniquely mappable ATAC-seq fragments per sample.

It was recently suggested that old cells have a globally lower nucleosomal occupancy that could result in permissive transcription and genomic instability [10]. Since that study was limited to very old cells, we used ATAC-seq data to look for changes in nucleosomal occupancy as a function of time and in different genetic backgrounds. Nucleosomes protect DNA from transposition by Tn5, resulting in depletion of Tn5 insertions in a region of about 140 bases around the centers of chemically mapped nucleosomes (Fig. 4A) [62]. These regions appeared to become more accessible in a sample from the old cells, consistent with global loss of nucleosomes. We noticed, however, that ATAC-seq samples from the old cells contained a higher level of background insertions. For instance, mid-gene bodies that are normally inaccessible to ATAC-seq became uniformly more accessible (Fig. 4B). This led us to examine an alternative hypothesis, that the non-specific background signal was uniformly higher in the samples from the old populations, leading to an apparent gain of accessibility in nucleosomal regions. Consistent with this hypothesis, the accessibility of canonically open nucleosome-free regions (NFRs) decreased with age (Fig. S11A). Indeed, since our data was normalized in a way that every sample has the same number of ATAC-seq insertions, increased insertions in the closed regions ‘compete’ with the insertions in the open regions, resulting in a seemingly decreasing accessibility of the open regions (Fig. S11A).

**Figure 4:**
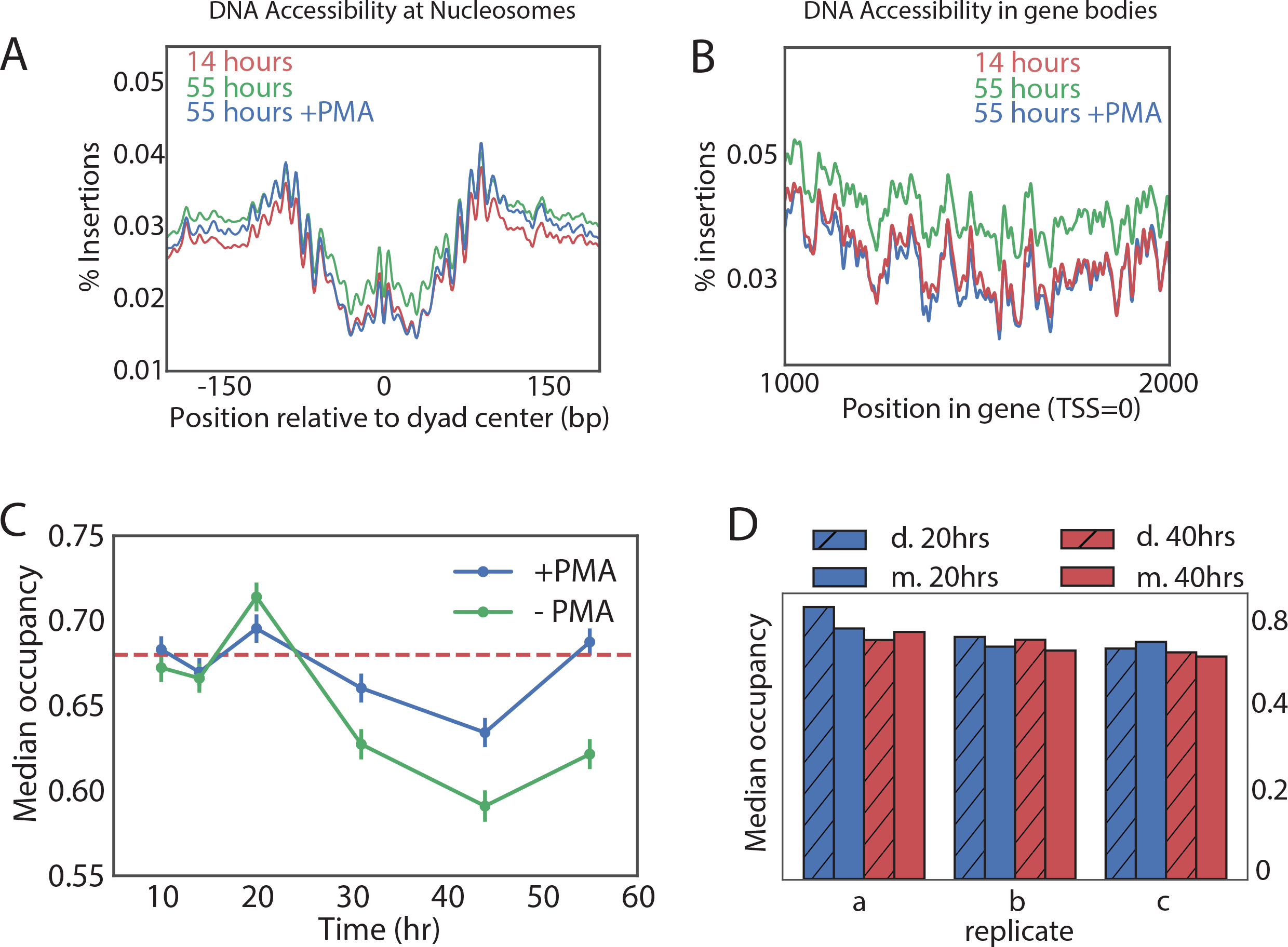
Nucleosome occupancy remains globally constant with aging. (A,B) Tn5 insertion density around well-positioned nucleosomes (A) and 1000 bases downstream of TSS (B) (Materials and methods) at 14 hours, 55 hours and 55 hours after treatment with PMA. (C) Median nucleosomal occupancy estimated by NucleoATAC on well positioned nucleosomes in open chromatin at different time points during time course. See Materials and methods for the method of estimation. (D). Same as (C) but comparing mothers and daughters during aging time course.

What can increase the background? Naked DNA from dead cells has been a prominent confounder in microbiological studies as extracellular DNA can remain stable for days [63]. DNA from the dead cells loses its chromatin structure, yielding a virtually uniform ATAC-seq signal (Fig. S12 top panel). Because the number of dead cells recovered (Fig. 1D, Table S1) increases with age, we hypothesized that the non-specific background in the ATAC-seq signal could rise because of the contamination by dead cells in old cultures. Since dead cells lose their membrane integrity, their DNA can be made inert to the ATAC-seq amplification by pre-treatment with intercalating agent propidium monoazide (PMA). PMA selectively enters dead cells and upon photo-activation binds covalently to DNA, strongly inhibiting its amplification in subsequent PCR reactions. Testing the efficacy of PMA treatment in yeast on heat-killed cells revealed that PMA can effectively remove the ‘dead cell’ signature from mixed samples (Fig. S12). We, therefore, compared ATAC-seq signal with and without pre-treatment of the sample with PMA.

Strikingly, addition of PMA decreased the accessibility of nucleosomal DNA in the old sample to the level observed in the young sample (Fig. 4A), and this PMA-dependent decrease in accessibility was specific to the old cells (>30 hours) (Fig. S11B). To verify that the average effect does not come from few exceptionally accessible nucleosomes, we binned nucleosomes into five accessibility bins and observed the same effect (Fig. S11C). Further, chromatin structure around promoters became virtually indistinguishable from that of the young cells and the non-specific background in the gene bodies decreased to the level of the young cells, consistent with a significant presence of the DNA from the dead cells in the old-cell samples.

To quantitatively measure the effect of age on nucleosomal occupancy we used NucleoATAC to measure nucleosomal occupancy around the well positioned nucleosomes in areas of open chromatin [61]. Indeed, without PMA treatment, average nucleosomal occupancy declined with age. This decline in occupancy, however, almost completely disappeared when the samples were treated with PMA (Fig. 4C). Also here, the effect of PMA was specific to the old cells, consistent with the increase in the dead cell proportion causing a seemingly declining occupancy (Fig. 4C). These findings were further supported by comparing occupancy of the PMA-treated daughter cell fraction (flow-through) to that of the middle aged and old mother cell fractions in three biological replicates (Fig. 4D). Also here, we observe that the nucleosomal occupancy remains constant, on average, with age (Fig. 4D).

We conclude that the global chromatin structure does not deteriorate significantly with age and previous observations to the contrary are likely caused by an increase in the proportion of the dead cells in the aging population.

### General and strain-specific age-dependent changes in chromatin accessibility

Although global chromatin structure did not change significantly between the young and the old cells, local changes in chromatin structure with age (Fig. S13A) were observed, suggesting that activity of multiple transcription factors and chromatin remodelers changes with age. To define age-dependent opening and closing of genomic regions, we divided the yeast genome into 100-bp long non-overlapping bins and performed a statistical test of age-dependent opening and closing (see *Materials and Methods*).

We first looked for common age-dependent chromatin changes between the four mutants. In general, at any significance cutoff more bins were closing than opening. We arbitrary set the significance cutoff at q-value of 10^−3^. At this q-value approximately 12% of bins were significantly opening or closing in all strains (Fig. 5A, S13B, Data S3). Bins located in gene bodies tended to preferentially open, while promoters tended to preferentially close. Origins of replications (ARS) strongly tended to lose accessibility (Fig. 5A, B, S13C, D). Comparing with other ATAC-seq datasets in yeast, we found that a similar closing of ARS elements occurs during the transition from the reductive to oxidative phase of metabolic cycle [64] (Fig. S13E). Since the reductive phase is characterized by a global decrease in DNA replication, our results suggests that DNA replication is inhibited in aging mothers.

**Figure 5:**
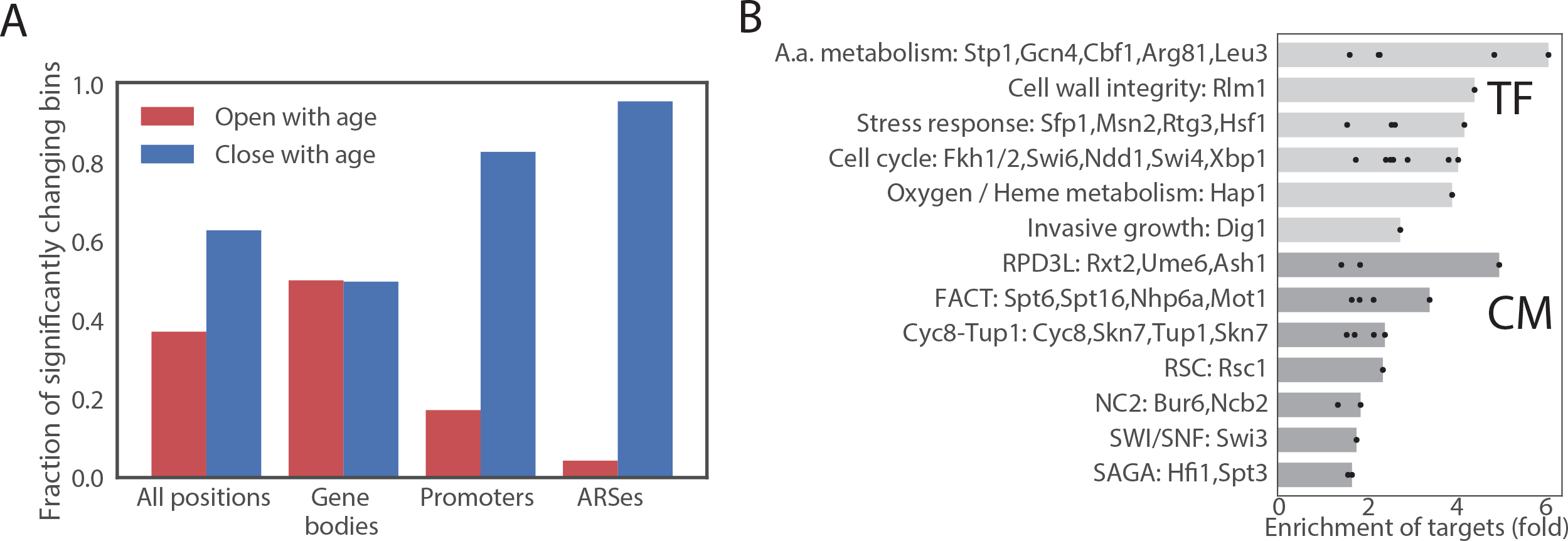
Age-dependent changes in chromatin accessibility. (A) Fraction of significantly changing bins that is opening or closing with age in each part of a genome. (B) Enrichments of genes regulated by particular transcription factors (TF) and chromatin modifiers (CM) among genes that contain bins that open with age. TFs/CMs are ordered in the order of increasing fold-enrichment and the bar marks the maximal fold-enrichment of a factor in the group.

We then asked if changes in chromatin accessibility reflect changes in activity of certain chromatin regulators or transcription factors through aging. We assigned each changing ATAC-seq bin to the closest gene and queried published datasets [65,66] if genes that contain changing bins are bound by a particular factor. No strong enrichment was found in bins that decrease in accessibility, but bins that increase in accessibility were highly enriched for binding of multiple transcription factors, including cell cycle regulators (Ndd1, Swi4/6, Fkh1/2 and Sbp1), and regulators of amino acid biogenesis, particularly Arg81, Leu3 and Gcn4 (Fig. 5B, Data S4). Notably, all enzymes in leucine biosynthesis superpathway and all enzymes in arginine biosynthesis pathway besides Arg2 and Arg7 gained accessibility with age.

To test if the age-dependency in chromatin accessibility differs between the mutants, we plotted the estimated age dependent slopes of bins in different genetic backgrounds. The aging slopes estimated in WT, *ubr2*∆ and *fob1*∆ were almost perfectly correlated with a slope of one, demonstrating that age-dependent changes in accessibility are very similar in these backgrounds. *Sir2∆* cells, however, behaved differently, with all slopes significantly higher than in the wild type, which we interpret as faster rate of aging in this background. Notably, however, although the slopes of changes were higher in *sir2*∆, overall, the slopes were highly correlated, suggesting that the overall pattern of age-dependent change in accessibility remains the same in this mutant strain (Fig. S13F).

### Accessibility and relative copy number at the rDNA locus is genotype dependent

Despite the remarkable similarity in aging ATAC-seq profiles across strains, we hypothesized that accessibility at the rDNA locus would be an important distinguishing feature as *SIR2* and *FOB1* are known modulators of rDNA stability. We reasoned that ATAC-seq insertional density at the rDNA locus would report on three distinct outcomes with respect to increasing instability: **1)** increased rDNA accessibility, **2)** increased rDNA copy number (extrachromosomal), and **3)** increased rDNA copy number (genomic). We found that with age, WT strains present with a ~2-fold increase in insertional density over the entire rDNA locus (Fig. 6, first panel). To determine whether the observed increase in ATAC-seq coverage at the rDNA locus was driven by increased accessibility **(1)** versus copy number expansion (**2 & 3**), we quantified relative rDNA copy number using qPCR with DNA from aged samples, and found that rDNA copy number increased with age (Fig. 6A). In fact, in comparison to insertional density, relative copy number increases to a larger extent with age, suggesting that the age-dependent increase in insertional density at the rDNA locus is largely being driven by copy number expansion, and average accessibility at the rDNA locus is decreasing (Fig. 6B). We observed that the increased ATAC-seq signal at the rDNA locus returned to normal levels in daughters born to old mothers, demonstrating an asymmetric division of signal. Together, these data argue strongly for **(3)** in that increasing ATAC-seq insertion density at the rDNA locus is reflective of greater numbers of asymmetrically retained ERC rDNA repeats with age.

**Figure 6:**
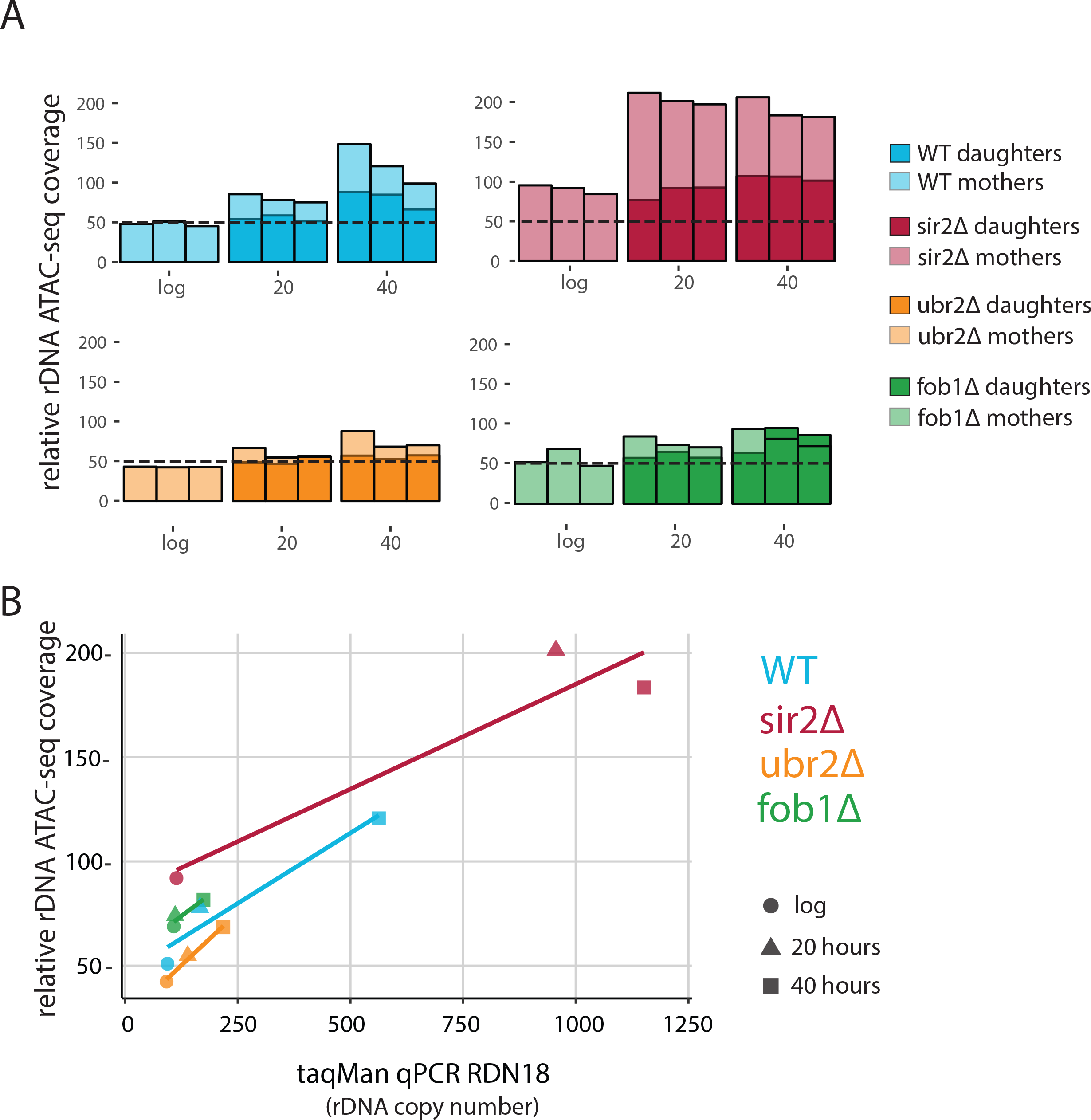
*fob1*∆ and *ubr2*∆ reduce age-dependent rDNA instability. (A) Percent relative (to non-repetitive genome) ATAC-Seq insertional density at the rDNA locus at increasing ages in both mother and daughter cells of WT and longevity mutants. Bars are ‘layered’ with data from daughter cells (opaque) displayed on top of data from mother cells (semi-transparent). (B) Comparing relative rDNA ATAC-Seq insertional density (y-axis) to relative rDNA copy number using TaqMan assay (x-axis) for multiple strains.

We looked at rDNA ATAC-seq coverage in *fob1∆* and *sir2∆* mutants, which repress and promote ERC formation, respectively. *sir2*∆ cells had slightly more (~20%) relative rDNA copy number than WT at young ages (Fig. 6A,B) but higher ATAC-seq coverage (2-fold), suggesting that during exponential growth, the difference between the strains is driven by bona fide differences in accessibility rather than by copy number. However, in looking at age related increases, relative rDNA copy number increases by ~10 fold and rDNA accessibility only increases twofold, suggesting that an initial open state leads to higher copy number with age, consistent with the well-documented ERC accumulation phenotype of *sir2*∆ mutants.

Surprisingly, like the *sir2*∆ strain, *fob1*∆ cells begin with ~15% higher relative rDNA copy number compared to WT during log phase, and have actually ~20% higher accessibility as measured by rDNA ATAC-seq than WT consistent with the role of Fob1p in Sir2p recruitment to the rDNA [67]. As expected, fob1∆ cells experience a much more modest increase in both ATAC-seq signal (1.4 fold vs 2.5 fold) and relative copy number (1.6-fold vs 6-fold) with age. Importantly, we note that if the source of our increased ATAC-seq rDNA coverage and estimated relative rDNA copy number are in fact from ERC build up, *fob1*∆ is only partially defective in ERC formation, consistent with previous work [68]. *Ubr2*∆ is a strain reported to modulate RLS independently of ERC formation [51]. In young cells, rDNA ATAC-seq signal and relative copy number for *ubr2*∆ was equivalent to WT. With age, however, the *ubr2*∆ cells displayed an attenuated rDNA phenotype compared to WT in both ATAC-seq (1.7 vs 2.5) and relative copy number (2.3 vs 6).

### *ubr2∆* and *fob1∆* strains exhibit stable nucleolar morphology with replicative age

In yeast cells, rDNA is localized, transcribed, processed and assembled into ribosomes within the nucleolar subcompartment of the nucleus [69]. It has previously been shown that the nucleoli of aged yeast cells grow large and fragment [70]. To compare our ATAC-seq measurements to a cell biological phenotype, we monitored nucleolar morphology for our mutants as a function of replicative age using the MAD platform. To do so, we aged cells from each strain that also harbored a fluorescent nucleolar reporter (a translational fusion of *NOP58* and mNeonGreen). Compared to young cells, all genotypes showed nucleolar enlargement (Fig. 7). However, the observed increases were clearly more pronounced in WT and *sir2∆* compared to *ubr2∆* and *fob1∆* (n ~ 50 cells for each strain and condition). Importantly, these results strongly support our conclusion that the changes we observed in ATAC-seq density at the rDNA locus with age correlate with a known aging phenotype related to rDNA destabilization. Furthermore, these data are consistent with the finding that like *fob1∆*, *ubr2∆* exhibits an attenuated decline in rDNA organization, a surprising result given that *UBR2* and upregulation of proteasome activity have been reported as independent from the *SIR2-FOB1* rDNA/ERC aging pathway [51].

**Figure 7:**
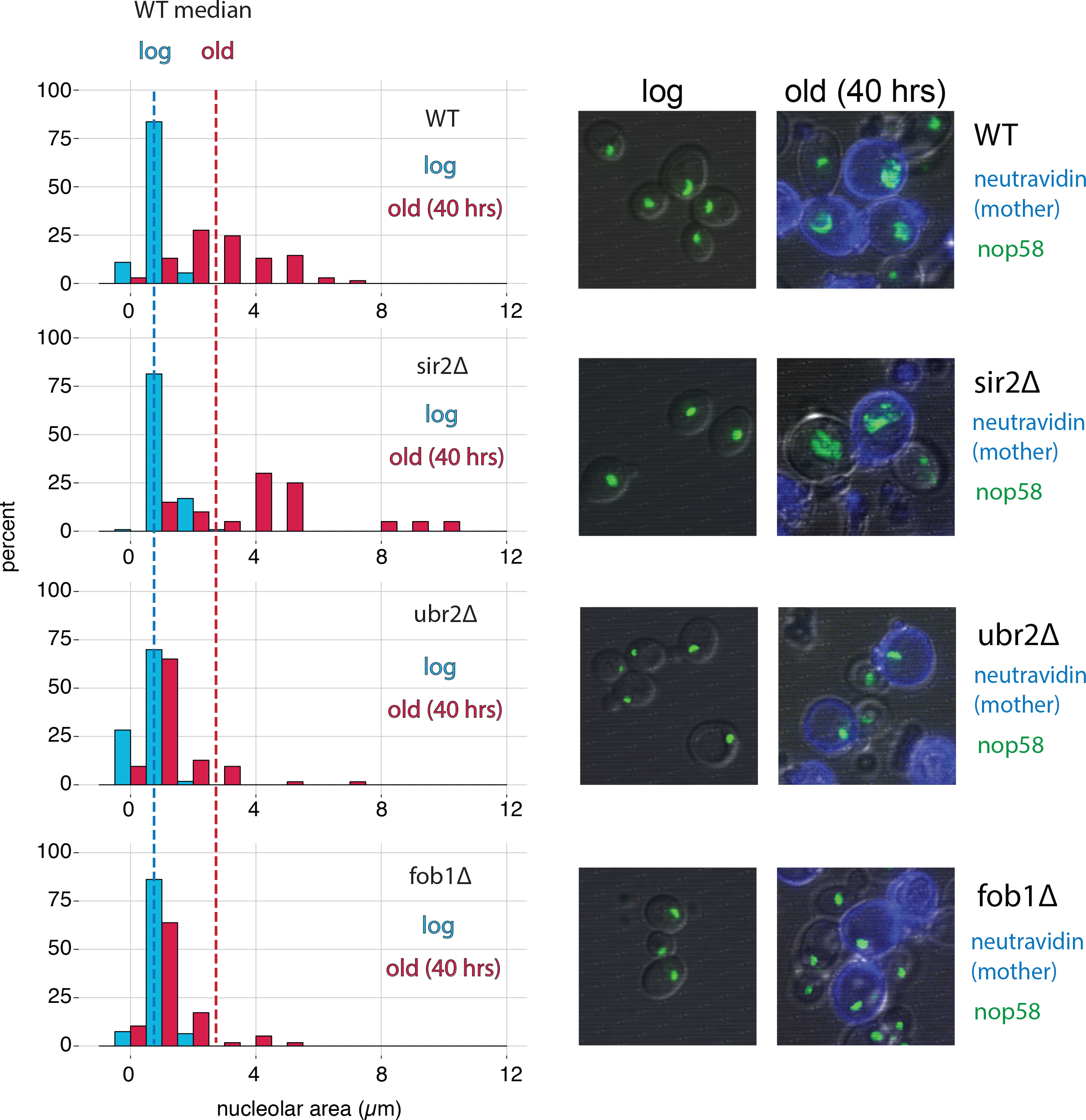
*fob1*∆ and *ubr2*∆ reduce age-dependent nucleolar fragmentation. (left) Nucleolar sizes in young (log) and old (40 hours) mother cells (displayed in blue and red, respectively) across multiple genotypes. (right) Visualization of nucleoli from young and old cells across genotypes.

## Discussion

Although the study of RLS in *Saccharomyces cerevisiae* has proven an informative model for studying how genetic and environmental factors impact cellular aging, the field has not been fully able to access the “omic” revolution. Significant technical challenges have impeded the collection of adequate numbers of viable aged cells for genomic or biochemical assays, much less aged cells across multiple environments and genotype. The extant systematic genomic research in yeast aging has been restricted to a handful of strains under similar conditions, none of which are tied to classic longevity genotypes or conditions (i.e., caloric restriction). Here, in order to fully leverage the environmental and genetic flexibility of yeast as an RLS model, we introduce the MAD platform as an effective methodology for aging large numbers of yeast cells in a wide variety of genetic background and environments.

We found in our analysis of the transcriptional response to aging that by far the dominant feature is the activation of the ESR, characterized by the induction of stress response factors and a repression of biogenesis programs that is highly correlated with growth rate. Although prior work has identified a connection between the ESR and aging [41], we show here the strength of that relationship as evidenced by the marked correlation between the rate that a gene changes with age, and the dependence of that gene’s expression on growth rate. We favor a model in which the concordance we observe suggests that much of the aging transcriptome is reflective of slowed growth in response to an as of yet unidentified insult. This model is consistent with reports of age-dependent reductions in the budding rates of single cells measured in microfluidics devices [71].

To highlight potential candidates responsible for age-induced ESR, we isolated stress-independent aging signatures by selecting genes that are not annotated as canonical ESR genes and whose expression is minimally dependent on growth rate (Fig. 3). Analysis of these genes revealed an early induction of aberrant transcription of the NTS region of the rDNA locus, transposon up-regulation, an oxidative stress response, a clear increase in the expression of SRP clients, and a cell wall integrity/ER stress signature that is independent of the ESR and GRS. Notably, we found evidence for all of these signatures in our mutant panel, albeit a much-attenuated transposon induction in the *sir2*∆ strain. Determining exactly how each of these events are interdependent versus distinct, and which might be causative, will require further study, presumably at the single cell level.

Previously, it has been reported that sub-telomeric regions are fast-evolving and the genes therein are enriched for being related to stress, cell surface properties, and carbohydrate metabolism, all of which are enriched functional annotations in our pan-aging gene expression signature [72]. Those authors speculated that the high evolvability of subtelomeric regions permits rapid adaptation to novel stresses [72]. From our analysis, close proximity to a telomere also increases the likelihood of age- and stress-dependent gene induction (Fig. S10). Whether this telomere position effect is due to a catch-all stress response that manifests with age, or perhaps an aging-specific redistribution of sirtuin proteins, remains an open question.

It has previously been reported that histone mRNAs and protein decline with replicative age in yeast [10,38]. This observation in tandem with a systematic measurement of nucleosome occupancy have accreted into a model wherein the reductions in histone protein lower the number of functional nucleosomes. The loss of nucleosomes is then purported to result in the erosion of appropriate transcriptional control which manifests as a universal up-regulation of all transcripts. Combining a modified ATAC-Seq protocol with MAD-purified old cells, we found that a global increase in DNA accessibility is likely not a general feature of aged cells. Importantly, we only observed a measurable increase in accessibility and decrease of occupancy at nucleosomes *without* the addition of the chemical agent PMA, which renders genetic input from dead cells inert. These results argue that previous reports of global nucleosomal loss in aging yeast cells may have been largely based on DNA from contaminating dead or dying cells.

We detected statistically significant changes in chromatin accessibility in ~12% of the genome. Strikingly, virtually all origins of replication are becoming less accessible as a function of age. Through comparison to other ATAC-seq datasets, we hypothesize, that in an aging yeast population, smaller fraction of cells are in S-phase and are thus are depleted for active replication, resulting in the observed declines at origins of replication. This is consistent with decreased expression of histones. As was the case with gene expression, the aging chromatin landscape was remarkably similar across our panel of mutants.

Changes in the rDNA stability with age manifest as ERC formation, which may be a common cause of replicative senescence in *S. cerevisiae* across numerous strain backgrounds and environments. Using our ATAC-seq data, we found, as expected, WT cells experience an increase in copy number of rDNA repeats that are asymmetrically retained. Likewise, we found an exaggerated and attenuated phenotype for *sir2*∆ and *fob1*∆ cells, respectively. We also found that the aged *ubr2*∆ rDNA profile phenocopies that of *fob1*∆, an unexpected result given that *UBR2* is believed to confer longevity through proteasome activation independently of *FOB1* and *SIR2*. Moreover, we verified this genomic observation using mutant strains with a genetically encoded, fluorescently tagged Nop58p to monitor nucleolar size and morphology as a readout for rDNA stability. Overall, these results are consistent with the known roles of *SIR2* and *FOB1* and provide provocative evidence that *UBR2* is either directly involved in ERC formation/rDNA stability, or that reduction in severity of rDNA aging phenotypes is a common feature of long-lived strains.

In the future, our expectation is that combining the MAD platform with single-cell analysis could be especially informative. MAD provides the means to obtain aged cells, which could then be used for single-cell sequencing and optical methods; indeed, single-cell RNA-seq could facilitate the identification of distinct aging trajectories where subtle causative changes can be distinguished from unsubtle stress responses [73,74], making it easier to separate cause from effect. In conclusion, by enabling the collection of multiple data modalities at different ages, genotypes, and environments, the MAD platform provides a powerful new paradigm for systems-level studies of cellular aging.

## Materials & Methods

### Cell preparation for aging in ministats

Cultures of desired strains were grown overnight at 30°C with shaking (180-200 rotations per minute [RPM]) and harvested at OD _600_< 0.2 by next morning (early log phase). Upon cells reaching the desired OD _600_, EZ-Link™ Sulfo-NHS-LC-LC-Biotin (ThermoFisher cat# 21338) was removed from −20°C freezer and equilibrated to room temperature during cell washing and preparation. For one ministat, 4 OD _600_ units of cells were washed twice in PBS + 0.25% PEG3350 at 1500g for 5 minutes at room temperature. During the second wash the NHS-LC-LC-Biotin was weighed out (0.5 mg per OD unit of cells) and re-suspended in 0.5 mL of PBS. Final washed cell pellets were resuspended in 0.5 mL of PBS and combined with NHS-LC-LC-Biotin in PBS for a final reaction volume of 1mL. The labeling reaction was carried out for 30 minutes at room temperature with rotation followed by a 2X wash in PBS + 0.25% as above. Biotin-labeled cells were resuspended in growth media and grown under normal liquid culture conditions (30°C with shaking at 180-200 RPM) for 10-14 hours. At time of seeding the number of labeled cells was determined using a Beckman Coulter Counter. In general, 4 OD _600_ units of cells produces between 130 and 170 million cells with variation arising from differences in strain size and cell loss during washing. Cultures were seeded such that the cell density at time of collection would not exceed an OD_600_ of 0.3.

Prior to cell collection, magnetic streptavidin beads (Dynabeads™ MyOne™ Streptavidin C1 ThermoFisher cat# 65001) were removed from 4°C storage, washed twice with 1 mL of growth media, and re-suspended in 20 mL of growth media. Following the post-biotin labeling phase of growth, cultures were passed over a 0.2 µm filter and re-suspended in 20 mL of growth media.

The number of beads used was 0.9 µL of beads per 1 million *labeled cells* (as measured previously with the Coulter counter [*e.g*., 135 µL of beads would be used for 150 labeled cells]). Well-mixed cells from the overnight culture (original mothers + all progeny) were collected for assays (RNA-seq, ATAC-seq, etc.) as the “log phase” sample.

Beading reactions were carried out by combining 20 mL of washed beads with 20 mL of re-suspended cells in a 50 mL falcon tube and rotating at room temperature for 15 minutes. Beaded mother cells were collected by placing the beading reaction on magnet (DynaMag™-50 Magnet ThermoFisher Cat#12302D) for 5 minutes. Supernatant (daughter cells) were carefully removed and discarded. Remaining beaded mother cells were washed 2X in 40 ml of growth media and re-suspended in a final volume of 1 to 2 mL of growth media. Beading efficiency was checked and cells counted under microscope with hemocytometer. Generally, beaded cells have between 1-10 beads per cell with an average of ~5 and recovery is usually around 75-90% of seeded mothers. Before loading onto ministat a sample of the beaded mothers was collected for assays.

For ministat loading, ministats were removed from magnets, the air pumps were set to lowest setting, and the effluent port was pulled far above media level. Beaded mothers were loaded through lauer-lock cap needle entry port using a 1000 µL micropipette. The loaded ministat was placed on magnet, and cells were allowed to bind for 10 minutes before restarting of pumps (60-75 RPMs) and air flow (25-33% of max output). Effluent needle was lowered to ~25 mm above the top ring magnet.

### Aging of cells in ministats

Loaded cells were grown on ministats for desired length of time. For the dense time course, 5 ministats were loaded and an entire ministat worth of cells was collected per time point. For short mutant strain time course, cells were harvested by removing ministat from magnet (pumps off, air low, and effluent needle up) and waiting 5 minutes from mothers to disperse. Sampling of cells was done through the loading port luer-lock syringe.

### Harvesting of daughter cells

Effluent tubing was diverted to a clean 50 mL Falcon tube to collect daughter cells. 5 million cells were spun down and frozen in liquid nitrogen for RNA, DNA preps or used immediately for ATAC-seq or buda scar counting as needed.

### Harvesting and washing of aged mother cells

Following removal of the ministat cap, tubing, and bubble cage (Fig. S14), the supernatant was removed using a 25 mL pipette, being careful not to disturb the magnetically-bound cells on the vessel wall. With the vessel removed from the magnet, beaded cells were resuspended in media and placed back on the magnet for 5-10 minutes. This was repeated for a minimum of 4 washes or until the cells were of desired purity as assessed by microscopy. Final beaded cells were resuspended in 1 mL of media and 5E6 cells were spun down and frozen in liquid nitrogen for RNA, DNA preps or used immediately for ATAC-seq or bud scar counting as needed.

### Bud scar counting and viability staining

Cells were resuspended in PBS + 1M sorbitol + 4mM EDTA and stained with 10 µL of NeutrAvidin Protein, DyLight (ThermoFisher 405 cat# 22831) to identify biotinylated mother cells, 1 µL 5 mg/mL Wheat Germ Agglutinin, Alexa Fluor 488™ Conjugate (ThermoFisher cat# W11261) to determine replicative age, and 1 µL of 5mg/ml propidium iodide) to quantify replicative age. Cells were stained for 30 minutes at room temperature and then washed twice in PBS with 1M sorbitol before microscopy. Viability and Purity analysis was carried out exactly as in [9]. FACS analysis was carried out with FloJo.

### mRNA extraction

Frozen cell pellets were resuspended in 200 µL of Lysis buffer (10mM Tris [pH 8.0], 0.5% SDS, 10 mM EDTA). Following the addition of 200 µL Acid Phenol [pH 4.3], samples were vortexed for 30 seconds. Samples were incubated at 65°C for 1 hour in a Thermomixer with intermittent shaking (2000 RPM for 1 minute every 15 minutes). 400 µL of ethanol was then added and the RNA was purified using the Direct-zol^TM^RNA Miniprep Plus kit (Zymo research) according to the manufacturer’s protocol including the optional DNase digestion step. RNA integrity was confirmed using an Agilent Bioanalyzer.

### RNA-Seq Library Preparation

RNA-seq libraries were prepared first by removing rRNA with the Ribo-Zero Gold rRNA Removal Kit (Yeast) (Illumina Cat #MRZY1324 and eluted in 30 µL. After rRNA removal, RNA was bound to AGENCOURT RNACLEAN XP beads (Beckman coulter cat#A66514) using a 2X bead volume (60 µL) followed by washing and elution as per vendor’s protocol (except that RNA was eluted in Fragment, Prime, Finish Mix from the TruSeq Stranded mRNA Library Prep [Illumina cat# 20020595]). Subsequent steps were carried out as per the protocol for the TruSeq Stranded mRNA Library Prep. Libraries were indexed using either Illumina barcode kit cat#s 20020492, 20020493 or 20019792). Libraries were sequenced on a hISEQ4000 (with 150×150 paired-end reads).

### RNA-Seq Data Processing and analysis

RNA-seq data was quantified using Salmon-0.8.1 with three separate annotation indices:

1. open reading frames (ORFs) as defined by SGD
2. “complex transcriptome” index created by aggregating ORFs, their longest annotated UTRs + all non-canonical and non-coding transcripts pulled from SGD (references)
3. all features index composed of ORFs + non-coding transcripts (SGD all freatures) + genomic intergenic bins

Salmon was run using the “quant” command with the following settings:“libType”: “ISR”, “useVBOpt”: [], “numBootstraps”: “30”, and “incompatPrior”: “0”.

Salmon quantification was prepared and processed for differential expression analysis with the Wasabi package in R for input into Sleuth. Genes changing with respect to time (generation) as opposed to batch or strain were defined as significant using: model = ~generation*strain+(batch)+(0)” and using a p <0.05 for calculated beta values. For determining genes changing significantly across time (age) for each specific strain the model “~batch+strain+strain:generation” was specified. All test tables output from Sleuth included in Data S1 and Data S2.

Batch correction (for heatmap visualization only) was applied to the Salmon quantification output matrix with a pseudocount of 2 TPM with the COMBAT package in R [75]. Hierarchical clustering (average linkage) was performed on the batch-corrected values using the uncentered Pearson correlation as the distance metric.

Gene Ontology (GO) term enrichment analysis was performed via uploading query lists to the DAVID Bioinformatics Resources 6.8 website and running enrichment with the *S. Cerevisiae* species background.

### Live/Dead Cell prep for ATAC-Seq

WT cells (DBY12000) were grown in YPD to OD_600_ 0.2 - 0.4, spun down (1200 rcf / 4 min./ room temperature), resuspended at 10^8^ cells/mL in YPD and split into two 1 mL aliquots for “live” and “dead” cells. The “dead” cell aliquot was incubated at 50°C for 25 minutes in a thermomixer and 100 µL (10 million cells) was plated on YPD to test viability (only 2 CFUs grew). Live and dead cells were combined into 100 µL aliquots at the desired ratios. ATAC-Seq was performed as described below.

### ATAC-Seq Sample Preparation

5 million cells were spun down (1200 rcf / 4 min. / room temperature) and resuspended in 250 mL media. Propidium Monoazide (PMA, Qiagen 296015) was added to a concentration of 50 mM, the cells were incubated in the dark at room temperature for 10 min. and then illuminated in the BLU-V system (Qiagen 9002300) for 20 minutes at 4^°^C. Cells were spun down, washed twice with SB buffer (1M Sorbitol, 40 mM HEPES [pH7.5], 10 mM MgCl_2_) and then resuspended in 190 mL SB Buffer. 10 µL of 10 mg/mL Zymolyase-100T was added and the cells were incubated at 30^°^C for 30 min. with light shaking (600 RPM) in a thermomixer (Eppendorf). Spheroplasts were spun down (1500 rcf / 5 min. / room temperature) and washed twice with SB Buffer, resuspended in 50 mL Tagmentation Mix (25 mL Nextera Tagment DNA Buffer, 22.5 mL H_2_O, 2.5 mL Nextera Tagment DNA Enzyme I) and incubated at 37^°^C for 30min. DNA was purified over DNA Clean & Concentrator^TM^-5 kit (Zymo Research) following the manufacturer’s protocol, eluted in 11ml H_2_O and stored at −20^°^C until ready for PCR.

PCR reactions were set up using Nextera Index i5 and i7 series PCR primers (Illumina). Mixed 25ml NEBNext Hi-Fidelity 2x PCR Master Mix, 7.5ml H_2_O, 6.25ml i5 primer (10 mM), 6.25ml i7 primer (10 mM) and 5 mL tagmented DNA from above such that each sample has a unique barcoded primer pair. Ran PCR amplification (1 cycle: 72°C for 5 min.; 1 cycle: 98°C for 30 sec.; 8 cycles: 98°C for 10 sec., 63°C for 30 sec., 72°C for 1 min.; hold at 4°C). PCR reactions were cleaned up using the Agencourt AMPure XP system (Beckman Coulter) 1^st^ by negatively selecting against large DNA fragments using 0.4x volume of beads and then by positively selecting for the desired fragments using 1.5x volume of beads. The final solution was re-purified over DNA Clean & Concentrator ^TM^-5 columns to eliminate primers, eluted in 22 µL H_2_O and analyzed on the BioAnalyzer.

### ATAC-Seq Data Analysis

#### Processing of raw sequencing data

Raw ATAC-seq data was processed as described in [76]. Briefly, first, the raw paired end reads were aligned to the Saccharomyces cerevisiae genome (sacCer3, Release 64) using bwa mem version 0.7.12 with default parameters. Second, the alignments were filtered requiring that both reads in the pair are mapped, the mapping quality is greater than or equal to 30, the reads are mapping concordantly and the direction of the mapping is F1R2. Third, we converted the alignments into track of number of insertions at each position in the genome. For every aligned fragment the insertion locations are calculated by shifting the fragment ends of reads that align to ‘+’ strand by 4 nucleotides and shifting the fragment ends of reads that align to ‘-’ strand by 5 nucleotides in the 3’ direction to reflect the distance to the center of the transposase binding site [60]. To normalize for uneven sequencing depth and different amounts of rDNA in the samples, insertion counts were normalized such that the total number of insertions from unique non-rDNA, nonmitochondrial reads is constant in each sample.

#### Annotations

Nucleosome positions were downloaded from [62], lifted over to sacCer3 assembly and only nucleosomes with mapping score greater than 2 were used in the metagene plots. For metagene plots around promoters we defined the TSSs for each yeast gene my merging the TSS data from [77] (TIF-seq) and [78] (TSS data). If the TSS for a gene was not reported by these two papers, we used the beginning of ORF as the TSS. TF and chromatin modifier binding data were from [65,66].

#### NucleoATAC

Baseline estimation of the nucleosome occupancy from ATAC-seq data could be confounded by noise in estimation of nucleosome free and nucleosomal fraction of fragments between samples. Therefore we generated a fragment size distribution that was an average fragment size distribution of all samples and ran the NucleoATAC pipeline using a constant fragment size distribution using –sizes parameter. Since occupancy estimation was sensitive to the depth of the sample, we downsampled each sample to 25 million uniquely mapped non-ribosomal reads. Finally, to limit differences due to differences in nucleosomal calling between samples, we called occupancy only on the nucleosomes from Brogaard et al that are well positioned (score > 2), located in the open chromatin part of the genome (as defined by Schep et al) and have a median occupancy of at least 0.5.

#### Differential accessibility

Aging slope for accessibilities was calculated as follows. We first binned the genome into non-overlapping 100 bp bins and counted number of insertions in each bin. We then performed quantile normalization for each sample separately to calculate normalization factors and supplied gene specific normalization factors as described in http://bioconductor.org/packages/release/bioc/vignettes/DESeq2/inst/doc/DESeq2.html#sample-gene-dependent-normalization-factors to DESeq2 pipeline [79]. The aging slope was determined on all samples using the following model: “~ replicate + strain + ave_budscars” where ave_budscars were the average number of budscars for that time point and we looked at the bins with q-value of non-zero slope of less than 10^−6^. To identify bins that were specifically opening or closing in certain strains, we used the following model “replicate + strain + ave_budscars + strain:ave_budscars”.

#### Enrichments of regulators

We assigned each genomic bin to the gene whose gene body it intersects or to the promoter (defined as up to 400 bp upstream of TSS) of the gene that intersects. The lists of age-opening and age-closing genes were generated by looking at a genes that contained at least one opening and closing bin (with q-value < 10^−6^) respectively. We then intersected these lists with the list of genes that bind to transcription factor or chromatin modifier and performed a hypergeometric test to evaluate enrichment.

### Strain Construction

Strains containing single-gene deletions were constructed using the standard PCR-mediated gene disruption method [80]. NOP58-3xmNeonGreen fusion proteins were constructed in WT, *sir2∆, ubr2∆* and *fob1∆* strains by PCR amplifying a 3xmNeonGreen/natMX cassette (from pKT127-3xNeonGreen-NatMX) using primers with homology to the NOP58 C-terminus and 3’UTR. All strains are S288c with a repaired *HAP1* allele [81]. A full list of strains used in this study can be found in Table S3.

### Nucleolar Quantification

Strains were grown in ministat media to mid-log phase (OD_600_ of 0.2 - 0.4), labelled and beaded according to the standard ministat protocol. Cells were harvested at log phase (pre-biotinylation), “young” (purified after initial 20 hours of outgrowth, before loading onto ministat) and “old” (after 23 hours of growth on the ministat, 43 hours of growth). Cells were stained with NeutrAvidin Protein, DyLight 405 (ThermoFisher cat# 22831) and Wheat Germ Agglutinin, Alexa Fluor™ 594 (ThermoFisher cat # W11262) to label the original biotinylated mothers and bud scars respectively.

Approximately 50,000 cells/well were loaded onto a Matriplate 384-Well glass bottom microwell plate (Brooks Life Science Systems cat # MGB101-1-1-2-LG-L) which had been pre-treated for 10 minutes with 1 mg/mL concavalin A. Cells were imaged at 1000x magnification on a VT-iSIM microscope (BioVision Technologies).

Images were analyzed in ImageJ and nucleolar area was determined by first creating a mask outline around original (biotinylated, Neutravidin-405-stained) cells. This mask was overlayed onto the mNeonGreen image, the threshold of the green areas was set and the analyze particles function was applied to calculate the nucleolar areas within each original cell. If the nucleolus was fragmented, the area was recorded as the sum of all fragments within that cell. Nucleoli outside of cells or within non-neutravidin-405 stained cells were not included in the analyses.

### qPCR analysis of rDNA

PCR primers and probes were designed against each target sequence using the online PrimerQuest Tool (IDT-Integrated DNA Technologies) and ordered as PrimeTime QPCR Assays (primer and probe mix) with probes containing a 5’ FAM reporter and ZEN / Iowa Black FQ double quencher. Each amplicon was also synthesized and cloned into a standard vector to generated copy # standard curves. Total DNA was isolated from frozen yeast pellets (5 million cells frozen in liquid nitrogen) via standard Zymolyase digestion/ Buffered Phenol:Chlorofom:Isoamyl (25:24:1, pH 8.0) extraction / EtOH precipitation methods. Each amplicon plasmid was diluted to generate a standard curve ranging from 10 to 781,250 copies/well. qPCR was run on 250 pg of total DNA as well as the standard curve using IDT’s PrimeTime Gene Expression 2x Master Mix in 384-well format on the Applied Biosystems™ ViiA™ 7 system. Data was normalized to *ACT1* copy number to obtain relative copy number per genome.

---

**Supplementary Figure 1: Enriched GO terms in the aging transcriptome.** (A) Volcano plot of ~2900 significantly changing transcripts (ORF + non-canonical transcripts) with age as modeled using Sleuth. The Non-Transcribed Sequence (NTS) of the rDNA locus is one of the largest measured increases (purple).

**Supplementary Figure 2: The effect of abundance on gene expression with age.** The aging slope for all transcripts was plotted against the mean natural log of sequence read counts across all samples. Each transcript (dot) colored by the q-value. In general low abundance transcripts are more likely to significantly increase rather than decrease with age.

**Supplementary Figure 3: Batch effects drive the age-dependent universal expression increase observed in Hu *et al*. 2014.** (A) Exact figure from Hu *et al* 2014 of spike-normalized RNA-seq data showing that in absolute terms, all transcripts increase with age. Replicate experiments were averaged. (B) Spike normalized data from Hu *et al* 2014 split into replicate experiments, “A” and “B”, and row normalized. Data from old cells from batch “A” are highlighted in red to point out that the majority of the universal up-regulation arises from this batch. (C) Re-analysis of the Hu *et al* raw data revealed exceptionally low percent reads mapping to the genome in general that was exacerbated in batch A. (D) Re-analysis of the spike-normalized data with batch “A” removed reveals that rather than a universal up-regulation, many genes change in both directions for this batch.

**Supplementary Figure 4: Characterization of mutant panel.** (A) Doubling times of each mutant and WT in standard minimal media and in ministat media (SD + 2% mannose + DAPA [no biotin]). (B) Verification of proteasome upregulation in the *ubr2∆* background. Gene expression changes for proteasome components in each mutant strain normalized to WT log phase. (C) Verification of deletions.

**Supplementary Figure 5: Quantifying the viability and purity of aged mother samples using flow cytometry.** Example of flow cytometric analysis of aged yeast cells.

**Supplementary Figure 6: Bud scar counts for MAD collected aged mutants.** (A) Bud scar counts for all yeast mothers cells at each timepoint during triplicate time courses. (B) The same data for replicate number 2 subset by viability.

**Supplementary Figure 7: Strain specific changes in the aging transcriptome of mutant panelhobb.** (A) Volcano plot displaying genes that change significantly with age in *sir2∆* compared to WT. Color indicates directionality – blue =decrease, red=increase (left panel). Scatter plot of aging slopes from WT (y -axis) versus *sir2∆* (x -axis) with colored points from the volcano plot colored similarly here. Diagonal line in black, regression line in blue (right panel). (B) same as in A, but for *ubr2∆* (C) same as in A, but for *fob1∆*.

**Supplementary Figure 8: SRP clients increase in expression as a function of decreasing SRP components.** For each perturbation in the Holstege data set, the average change in SRP components was plotted against the average change for all SRP clients. The negative correlation reveals that as SRP components go down in expression, SRP targets go up in expression.

**Supplementary Figure 9. *SCR1* expression increases with age in the dense WT time course.** For each timepoint, *SCR1* expression in Transcripts per Million (TPM) was plotted.

**Supplementary Figure 10: Telomere proximal genes increase with both replicative age and stress** (A) The Loess fit for aging slope plotted against gene position (relative to telomeres) for all genes from each longevity mutant. (B) The Loess fit for growth rate slope plotted against gene position (relative to telomeres) for all genes measured for each longevity mutant.

**Supplementary Figure 11: PMA treatment rescues young cell ATAC-seq profile at transcriptional start sites in old cell populations.** (A) Metagene plot of ATAC-seq insertion density for genes aligned at the transcription start sites for young cells (red), old cells (green) and old cells with PMA treated (blue). (B) Effect of PMA is strongest at nucleosomes with low accessibility. We binned nucleosomes into five bins according to their average occupancy and observed that the nucleosomes in the low-accessible regions are affected the most consistent with addition of a constant background to the accessibility signal. (C) Insertion density around well-positioned nucleosomes before and after treatment with PMA at various times along aging time course.

**Supplementary Figure 12: Efficacy of PMA in removing the ATAC-seq signature of heat-killed cells from mixed populations.** Depth-normalized ATAC-seq coverage was plotted at specific locus to illustrate the lack of structure in ATAC-seq data from heat-killed cells (top panel) and that treatment with PMA effectively rescues live cell signal from a mixed live/dead population over a range of concentrations (panels 2-6).

**Supplementary Figure 13:** (A) Multiple local changes in chromatin during aging. We show a hierarchically clustered heatmap of bins that significantly change with age (newborn cells at 20 hrs (NB), 20 hrs of aging and 40 hrs of aging) in all profiled strains (see Material and Methods). (B) Number of genomic bins significantly changing accessibility with age at each threshold of significance. (C) Distributions of fold changes of significantly changing bins that are in promoters, gene bodies, origins of replications and everywhere. Note that almost all significantly changing bins in ARSes (250) decrease in accessibility with age. (D) Differences in patterns of changes of accessibility between promoters, gene bodies and origins of replication can’t be explained by average initial accessibility of each genomic location type. We binned each type of genomic locations into bins according to accessibility in Log phase and asked what proportion of bins with this level of accessibility go up in each type of genomic location. Notably, at the same level of accessibility ARSes tend to close the strongest. (E) Closure of ARSes in metabolic cycle. Shown are changes in occupancy similar in transition from RB to OX phase of the metabolic cycle (data from [64]). Similar to (C). (F) Comparisons of aging slopes of genomic bins between mutants.

**Supplementary Figure 14: Images of the MAD platform.** (A) Side-view of a single MAD. The “Bubble Cage” enables aeration of the culture, and prevents large bubbles from dislodging beaded mother cells from the sidewall of the ministat. (B) View of four MADs.

**Table Captions**

**Table S1: Physiological parameters collected from aging time courses.**

**Table S2: Yeast transcript types.**

**Table S3: Strains used in this study.**

**Table S4: TaqMan primers and amplicons.**

